# The Quercetin and Quercetin Derivatives Interaction With Cyclooxygenase-1 and Cyclooxygenase-2

**DOI:** 10.1101/2020.12.05.413088

**Authors:** A.E. Manukyan, A.A. Hovhannisyan

## Abstract

The cyclooxygenase (COX) enzymes are tumor markers, the inhibition of which can be used in the prevention and therapy of carcinogenesis. It was found that COX-1 and COX-2 are considered as targets for tumor inhibition. In anticancer therapy, plant compounds are considered that can inhibit their activity. Modeling of the COX-1 and COX-2 enzymes was carried out on the basis of molecular models of three-dimensional structures from the PDB database [PDB ID: 3KK6, 5f19] RCSB. For docking analysis, 3D ligand models were created using MarvinSketch based on the PubChem database [CID: 5280343, 5281654]. *In silico* experiments, for the first time, revealed the possible interaction and inhibition of COX-1 and COX-2 by quercetin and quercetin derivatives. Aspirin and Celecoxib [CID: 2244, 2662] were taken to compare the results. Possible biological activities and possible side effects of the ligands have been identified. It is noteworthy that Celecoxib is not active on the studied cell lines, while quercetin and quercetin derivatives are more active than Aspirin.

## INTRODUCTION

The molecular mechanisms of drugs are the foundation on which can be built the therapy of many diseases accompanied by inflammatory processes. [1]. Molecules of various biochemical origins are involved in the regulation of inflammatory process. Eicosanoids are highly active regulators of cellular functions, have a short T1/ 2, therefore they have effects as “local hormones”, occupy one of the central positions in inflammatory processes: they include prostaglandins, thromboxanes and other substances. Excessive secretion of eicosanoids leads to a number of diseases. The main substrate for the synthesis of human eicosanoids is arachidonic acid [8]. The synthesis of arachidonic acid is carried out due to the “work” of varieties of the cytoplasmic enzyme phospholipase A2, which in turn is activated by proinflammatory signaling pathways. After the process of separating arachidonic acid from the phospholipid, it is released into the cytosol and converted into different eicosanoids depending on the cell types. The most famous is the classic type of eicosanoids, which are formed due to the activity of the enzymes cyclooxygenase-1 (COX-1) and cyclooxygenase-2 (COX-2). The COX enzyme catalyzes the first step in prostaglandin synthesis. The released arachidonic acid is converted by COX to prostaglandin (PGs) –PGG2, which is reduced to prostaglandin PGH2, and that into other eicosanoids (prostaglandins, thromboxanes, prostacyclins). PGs are cellular mediators of inflammation, the violation of biosynthesis of which can cause the development of severe pathological conditions: participation in the implementation of dysplastic and neoplastic processes (tumor growth), namely, in the suppression of apoptotic cell death, pathological neoangiogenesis and invasion, as well as mediation of immunosuppressive functions [7]. COX-1 is expressed in almost all mammalian tissues, and COX-2 is constitutive in some parts of the central nervous system. At the same time, COX-2 is rapidly activated in response to proinflammatory mediators and mitogenic stimulants [13]. A decrease in the recurrence rate of many tumors was noted with the use of non-steroidal anti-inflammatory drugs (NSAIDs) that inhibits COX-2, but all these compounds, together with inhibition of COX-2, also inhibit the activity of COX-1, which leads to a negative effect on the implementation physiological processes in the body. Thus, the search for non-toxic and effective COX-2 inhibitors continues to this day [7; 10]. Recently, researchers have been paying special attention to substances of natural origin that have antitumor activity and are capable of selectively inhibiting the activity of COX-2 [10]. Some of these compounds can be quercetin and quercetin derivatives. Numerous studies have revealed the anticarcinogenic effect of quercetin and quercetin derivatives, which can prevent endothelial dysfunction, the production of superoxide caused by angiotensin II, inhibit the proliferation of tumor cells, causing cell apoptosis [2; 6; 9].

The aim of this work is to study the probable interaction of quercetin and quercetin derivatives by molecular docking with COX-1 and COX-2, in comparison with Aspirin and Celecoxib [7; 10; 12].

## RESEARCH METHODS

The search for ligands and targets was carried out on the basis of verified articles from the Pubmed data bank. The structural formulas for the ligands were taken from PubChem [CID: 5280343] and constructed by MarvinSketch. All possible targets for each ligand were identified [13]. Repeats were removed from ∼ 650 targets and were selected 25 targets belonging to *H. sapiens*. Of the 25 selected targets were examined COX-1 and COX-2. Target modeling was carried out on the basis of molecular models of three-dimensional structures PDB RCSB [ID: 3KK6, 5F19]. Physicochemical parameters were determined using SwissADME (Table 1) [3]. Docking analysis carried out by using AutodockVina, a reliability analysis of which is provided by 40-fold repeatability for each ligand [11].

**Table 1.**

| Physico-chemical<br>parametres of the<br>ligands | Aspirin | Celecoxib | Quercetin | 3-Methylquerc<br>etin | Quercetin<br>3-O-rhamnoside |
| --- | --- | --- | --- | --- | --- |
| Formula | $C_9H_8O_4$ | $C_{17}H_{14}F_3N_3O_2S$ | $C_{15}H_{10}O_7$ | $C_{16}H_{12}O_7$ | $C_{21}H_{20}O_{11}$ |
| IUPAC<br>name | 2-acetyloxyb<br>enzoic acid | 4-[5-(4-methylphenyl<br>) -3-<br>(trifluoromet<br>hyl)pyrazol-1-yl]benzenesul<br>fonamide | 2-(3,4-dihydroxyphe<br>nyl)-3,5,7-trihydroxychr<br>omen-4-one | 2-(3,4-dihydroxyphenyl<br>) -5,7-dihydroxy-3-methoxychromen<br>-4-one | 2-(3,4-dihydroxyphen<br>yl)-5,7-dihydroxy-3-<br>[(3*S*,4*S*,5*S*,6*R*)-<br>3,4,5-trihydroxy-6-methyloxan-2-yl]oxychromen<br>-4-one |
| Num. rotatable<br>bonds | 3 | 4 | 1 | 2 | 3 |
| **H-acceptors;**<br>**donors** | 4;1 | 7;1 | 7; 5 | 7; 4 | 11; 7 |
| **Solubility** | 2.54 e<sup>0.2</sup> | 1.04 e<sup>-0.2</sup> | 2.11 e<sup>0.1</sup> | 4.07 e<sup>-0.3</sup> | 3,69 e |
| **GI absorbtion** | high | high | high | high | low |
| **BBB permeant** | yes | no | no | no | no |
| **log Kp (skin**<br>**permeation) sm/s** | -6.55 | -6.21 | -7.05 | -6.31 | - 8.42 |
| **Lipinski rule** | yes | yes | yes | yes | yes |
| **Ghose** | yes | no | yes | yes | yes |
| **Veber** | yes | yes | yes | yes | no |
| **Bioavailability**<br>**Score** | 0.56 | 0.55 | 0.55 | 0.55 | 0.17 |

## RESULTS AND DISCUSSION

For a detailed study of the ligands interaction with targets, docking analysis was performed with the detection of binding sites (Table 2) and energy characteristics (Tables 3, 4).

**Table 2.**

| **Target** | **Aspirin** | **Celecoxib** | **Quercetin** | **3-methyl-**<br>**quercetin** | **quercetin 3-**<br>**O-rhamnoside** |
| --- | --- | --- | --- | --- | --- |
| **COX-1** | S530; R120;<br>Y385; H90;<br>P156 | S530; R120;<br>Y385; H90;<br>F381 | Y355; V349;<br>Y385; R120 | Y355; V349;<br>Y385; R120,<br>H90 | Y355; V349;<br>Y385; R120,<br>H90 |
| **COX-2** | S530; R120;<br>Y385; V523;<br>H90; P156 | V523; R120;<br>Y355; Y385;<br>S530; H90 | P156; H39;<br>P153, V523,<br>R513 | G135;P156;<br>H39; V523,<br>P153, R513 | P156; H39;<br>V523, P153,<br>R513 |

**Table 3.**

| Ligand | COX-1 |  | COX-2 |  |
| --- | --- | --- | --- | --- |
| | Gibbs energy (kcal/m) | Binding constant $K_b$ ($10^4$) | Gibbs energy (kcal/m) | Binding constant $K_b$ ($10^4$) |
| Aspirin | -6.5±0.05 | 7,8 | -6.5±0.05 | 7,8 |
| Celecoxib | -9.1±0.05 | 11,2 | -8.8±0.05 | 10,8 |
| Quercetin | -9.4±0.05 | 11,4 | -9.7±0.05 | 11,7 |
| 3-methylquercetin | -8.5±0.05 | 10,5 | -9.7±0.05 | 11,7 |
| Quercetin 3-*O*-rhamnoside | -8.6±0.05 | 10,7 | -9.7±0.05 | 11,7 |

**Table 4.**

| Ligand | Glioma cells (Hs683) | Keratinocytes (HaCaT) |
| --- | --- | --- |
| Aspirin | 0.450 | 0.234 |
| Celecoxib | - | - |
| Quercetin | 0.458 | 0.205 |
| 3-methylquercetin | 0.399 | 0.177 |
| Quercetin 3-*O*-rhamnoside | 0.405 | 0.194 |

The active sites of COX-1 and COX-2 are similar - R120, Y355, E524, the folding of these amino acids leads to the formation of a pocket. Unlike COX-1, COX-2 replaced I523 with V523 and R513 with H513, which leads to the formation of an additional side pocket, which makes the active site of COX-2 more accessible, in addition, the last D helix curls up and facilitates binding to R120. The catalytic sites of COX-1 and COX-2 are also similar, and amino acid residues are formed by Y385 and S530.

It was found that when ligands interacted with COX-1, all bound to the active site Y385, R120, and also all ligands except quercetin bind to H90, forming hydrophilic bonds. Quercetin and quercetin derivatives bind to V349, and Celecoxib to F381, forming hydrophobic bonds.

In the case of COX-2, the ligands form hydrophilic bonds with V523, which is an additional pocket of the catalytic center. Aspirin and Celecoxib form hydrophilic bonds with R120 and Y385. Quercetin and quercetin derivatives, in contrast to Aspirin and celecoxib, interact with R513 and P156, which are involved in the organization of the catalytic center.

The data obtained indicate the possible binding of all ligands to targets.

For each ligand, the prediction of the cytotoxic effect of chemical compounds with non-transformed (HaCaT) and transformed (Hs683) cell lines was carried out, as a result of which the probable biological activities and possible side effects of the ligands were identified (Tables 3, 4).

It is noteworthy that Celecoxib is not active on the studied cell lines, while quercetin and quercetin derivatives are more active than Aspirin.

It follows from the results that quercetin and its derivatives are characterized by low cytotoxicity for untransformed cells, and the interaction with targets-COX-1 and COX-2 is stronger than with Aspirin and Celecoxib.

Author: Manukyan Amalya

Institute: Russian-Armenian University, Bioengineering, Bioinformatics and Molecular Biology

Street: Hovsep Emin 123 str.

City: Yerevan

Country: Armenia

## REFERENCES

1. Al-Salam S et al. Epstein-Barr virus infection correlates with the expression of COX-2, p16(INK4A) and p53 in classic Hodgkin lymphoma. Int J Clin Exp Pathol. – V. 2013; 6(12), P. 2765–77

2. Chakraborty A. et al. An efficient protocol for in vitro regeneration of Podophyllum hexandrum, a critically endangered medicinal plant. Indian Journal of Biotechnology 9(2):217–220 2010 V. 09(2). P. 217–220.

3. Daina A., Michielin O., Zoete V. SwissADME: a free web tool to evaluate pharmacokinetics, drug-likeness and medicinal chemistry friendliness of small molecules //Scientific Reports. – 2017 – V. 7 – P.42717.

4. Fantini M. et al. In vitro and in vivo antitumoral effects of combinations of polyphenols, or polyphenols and anticancer drugs: perspectives on cancer treatment //International journal of molecular sciences. – 2015 – V. 16(5). – P. 9241–9249.

5. Hashemzaei M. et al. Anticancer and apoptosis-inducing effects of quercetin in vitro and in vivo //Oncology reports. – 2017 – V. 38 – №. 2 – P. 819–828.

6. Khan F. et al. Molecular targets underlying the anticancer effects of quercetin: an update //Nutrients. – 2016 – V. 8 – №. 9 – P. 529

7. Leng J, Han C, Demetris AJ, Michalopoulos GK, Wu T-Cyclooxygenase-2 promotes hepatocellular carcinoma cell growth through Akt activation: evidence for Akt inhibition in celecoxib-induced apoptosis. Hepatology. 2003 – V. 38(3) – P.756–68.

8. Paul AG, Chandran B, Sharma-Walia N. Cyclooxygenase-2-prostaglandin E2-eicosanoid receptor inflammatory axis: a key player in Kaposi’s sarcoma-associated herpes virus associated malignancies. Transl Res. 2013 – V. 162(2) – P.p77–92

9. Raja S. B. et al. Differential cytotoxic activity of Quercetin on colonic cancer cells depends on ROS generation through COX-2 expression //Food and Chemical Toxicology - 2017 – V. 106– P. 92–106.

10. Ribeiro D., Freitas M., Tomé S., Silva A., Laufer S., Lima J., Fernandes E. Flavonoids Inhibit COX-1 and COX-2 Enzymes and Cytokine/Chemokine Production in Human Whole Blood. Inflammation 2014 – V. 38(2)

11. Trott O., Olson A. J. AutoDock Vina: improving the speed and accuracy of docking with a new scoring function, efficient optimization, and multithreading // Journal of computational chemistry. 2010 V. 31(2). – P. 455–461.

12. Xiao X., Shi D, Liu L, Wang J, Xie X, Kang T, Deng W. Quercetin suppresses cyclooxygenase-2 expression and angiogenesis through inactivation of P300 signaling. PLoS One. 2011; – V. 6(8)

13. Yao Z. J. et al. TargetNet: a web service for predicting potential drug–target interaction profiling via multsi-target SAR models //Journal of computer-aided molecular design. – 2016 – V. 30 – №. 5 – P. 413–424.

